# The Human Metabolome and Machine Learning Improves Predictions of the Post-Mortem Interval

**DOI:** 10.1101/2025.03.24.644974

**Authors:** Rasmus Magnusson, Carl Söderberg, Liam J. Ward, Jenny Arpe, Fredrik C. Kugelberg, Albert Elmsjö, Henrik Green, Elin Nyman

## Abstract

An accurate prediction of the time since death, known as the post-mortem interval (PMI), remains a critical research question in forensic and police investigations. Current methods, such as rectal temperature or vitreous potassium levels, only provide reliable PMI estimations up to 48-72 hours. In this study, we utilized metabolomic data from femoral whole blood samples of 4,876 individuals with known PMIs ranging from 1 to 67 days. We developed a neural network model that predicted PMI with a mean/median absolute error of 1.45/1.03 days in unseen test cases, outperforming six other machine learning architectures. To further highlight the biological signal, we performed pseudo time-series clustering of metabolic features used by the model, revealing 158 decreasing, 254 increasing, and 398 features with more complex patterns over the pseudo-time scale. Our findings also indicate that metabolomic data from approximately 256 individuals is sufficient to train a machine learning model for PMI prediction, making this approach widely applicable for researchers and forensic institutes worldwide.

## Introduction

Accurately determining the time of death or the time elapsed since a person died, also known as the postmortem interval (PMI), is of critical importance in forensic and police investigations. In general casework, knowledge of PMI can significantly improve the understanding of the circumstances surrounding the death of the deceased. An accurate estimate of PMI in a homicide case significantly impacts the ongoing investigation and the ability to reconstruct the chain of events that lead to death.

Despite considerable research [1]–[3], current best practices are unable to provide accurate and reliable estimates of PMI in a variety of scenarios. Typically, one or several of the following indicators are used to estimate PMI of a corpse: livor mortis (discoloration of the skin), rigor mortis (stiffening of the body), algor mortis (body cooling to surrounding temperature), and quantitative measurements of potassium concentration in vitreous humor. The accuracy of these methods varies. Generally, the evaluation of rigor mortis and livor mortis is subjective and can vary significantly between individuals [4].

The current gold standard methods for PMI estimation include measuring the rectal body temperature [3]. This method is effective until the body reaches the surrounding temperature [3], which typically occurs 24-48 hours post-mortem [3]. Another reliable method is measuring the potassium concentration in the vitreous humor of the eye, which increases proportionally with PMI [5]. This increase is linear for the first 24 to 48 hours, but after this period, the uncertainty of PMI estimate increases remarkably [5].

Past the early post-mortem time periods, methods are typically less precise and mostly cover substantially later PMI. Factors that can be used to predict PMI include insect activity [3], body decomposition [3], [6], [7], skeletonization, and bone bleaching. Indeed, there is a need for more precise methods for PMI determination that can be applied not only during the first 2-3 days post-mortem, but also in a longer postmortem time frame without sacrificing accuracy.

The human metabolome, which includes all endogenous substances of low molecular weight, has shown promising potential in PMI estimations when analyzing biochemical changes in body fluids and/or body tissues after death [8]–[14]. PMI has been identified as the main factor driving post-mortem metabolomic changes in various soft tissues and fluids [10]. However, current studies are conducted mainly using animal models, which can limit their application in forensic case work [9], [13].

Some studies have been conducted on human material. For example, studies have reported a correlation between PMI and the levels of metabolites such as hypoxanthine, choline, creatine, betaine, glutamate, and glycine in serum, vitreous humor, and aqueous humor [11] and threonine, tyrosine and lysine in muscle tissue [14]. In addition, taurine, glutamate, and aspartate in vitreous humor have also been associated with PMI [15]. However, there is still a lack of an accurate description of a prediction model using human data that would be suitable for forensic and police investigations.

In this study, we therefore used post-mortem metabolomic data collected from femoral whole blood samples from 4,876 individuals with known PMI, ranging from 1 to 67 days. We developed a neural network model that accurately predicts PMI with a mean absolute error (MAE) of 1.45 days and a median absolute error of 1.03 days. Furthermore, we explored various of-the-shelf supervised machine learning methods, finding that four of the six models demonstrated significant predictive power, although none surpassed the performance of the neural network model. These findings suggest that the metabolome can be effectively used to predict PMI, especially in the time frame of 2-10 days, a common time window in forensic investigations.

## Results

### A neural network model fills the gap of reliable PMI estimations in critical time frames

We used metabolomics data from 4,876 femural blood samples from forensic investigtion, with 2,305 selected features (see Methods). For these samples, 95% of PMI values were between 2 and 14 days, with the median PMI being 5 days. We randomly divided the samples into training, validation, and test sets in proportions 80% (3,907 profiles), 10% (471 profiles), and 10% (471 profiles), respectively (Fig. 1a).

**Figure 1:**
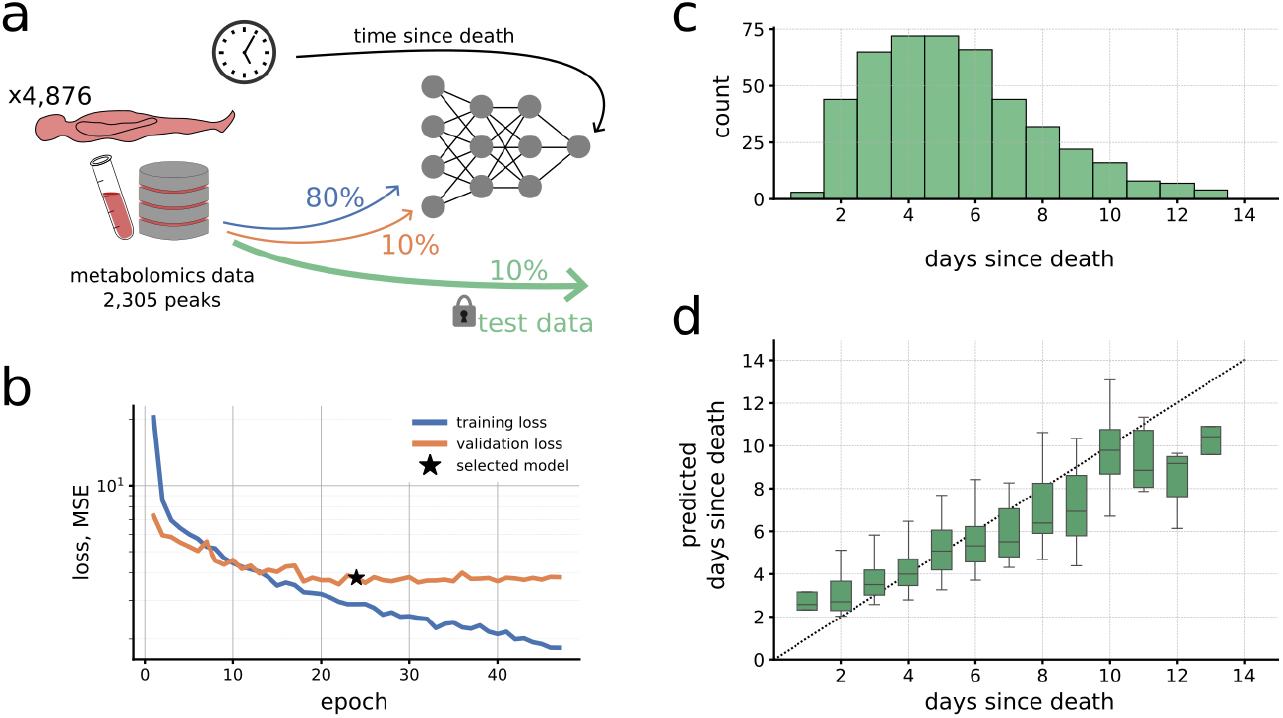
Study design and model performance. a) We used our compendium of 4,876 post-mortem metabolomic profiles to train a feed forward neural network (FFNN) regression model to predict the post-mortem interval (PMI). b) We trained the model with the best performing hyper-parameters using early-stopping with respect to the validation error. The training and validation losses are shown as a function of epochs. c) The test data typically had a PMI of 2-10 days, as shown in this histogram. The distribution was in line with that of the training and validation data. d) We found a mean absolute error of 1.45 days when predicting the PMI in the test data. Shown are the distributions of model estimates as a function of the actual PMI.

Given the number of samples, we chose to apply a feed forward neural network (FFNN) regression model to predict PMI. We first performed a hyperparameter optimization with respect to the model performance on the validation data. In detail, we tested 30 FFNN models with 1-4 hidden layers, each containing 32 to 512 hidden nodes. In addition, each layer had a dropout rate of between 0.05 and 0.5. The model always terminated in an additional single output node, and all activation functions used the rectified linear unit (ReLU). The models were implemented with a learning rate between 1e-4 and 1e-2. Furthermore, we explored whether incorporating feature selection as the initial step would improve model performance by implementing a custom-built attention mechanism as the first layer in the optimization process. The model selection included an early stopping with respect to the validation error, with a patience of 25 epochs.

We observed a noticeable robustness in model performance with respect to hyperparameter design. All tested model hyperparameter settings and their respective performances can be found in Supplementary Table (Table S1). Interestingly, we observed a clear trend where the performance was clearly stratified by the implementation of the custom-built attention layer. The 18 best-performing models all had such an attention layer, whereas the nine worst-performing models did not. The overall best-performing model (Fig. 1b) contained an instance of the attention layer, a dense layer with 288 hidden nodes with a dropout rate of 0.15, and an output layer. The optimal learning rate for this model was set to 0.005.

Having selected and applied the FFNN regression model to the test data and compared the predicted PMI values with the actual PMI values, we found the model to accurately predict PMI values, with a mean absolute error of 1.45 days (Fig. 1c). The median absolute error was 1.03 days. Given that some of the actual PMI values had a resolution of 48 hours (n=985), with a typical uncertainty of less than 24 hours, the relative error of the predictions was comparable to the underlying uncertainty of the training data.

### A compendium of supervised machine learning methods shows significant information embedded in the data

Neural networks are known for their ability to learn from complex, non-linear patterns in data, and for their requirement of large datasets to do so. We continued the study by analyzing whether less complex alternative machine learning models could also accurately predict PMI. To this end, we built a compendium of models and fitted them to the training data. The model types included in this compendium were a ridge regressor, a least absolute shrinkage and selection operator (LASSO), a support vector regression (SVR), a random forest regressor, a gradient boosting regressor, and a K-nearest neighbor regressor (k-NN). We again trained these models using 80% of the data (3,907 profiles) and tested them using 10% of the data (471 profiles). We observed clear predictive power across five of the six models, although the FFNN had the lowest mean absolute error (Figure 2).

**Figure 2:**
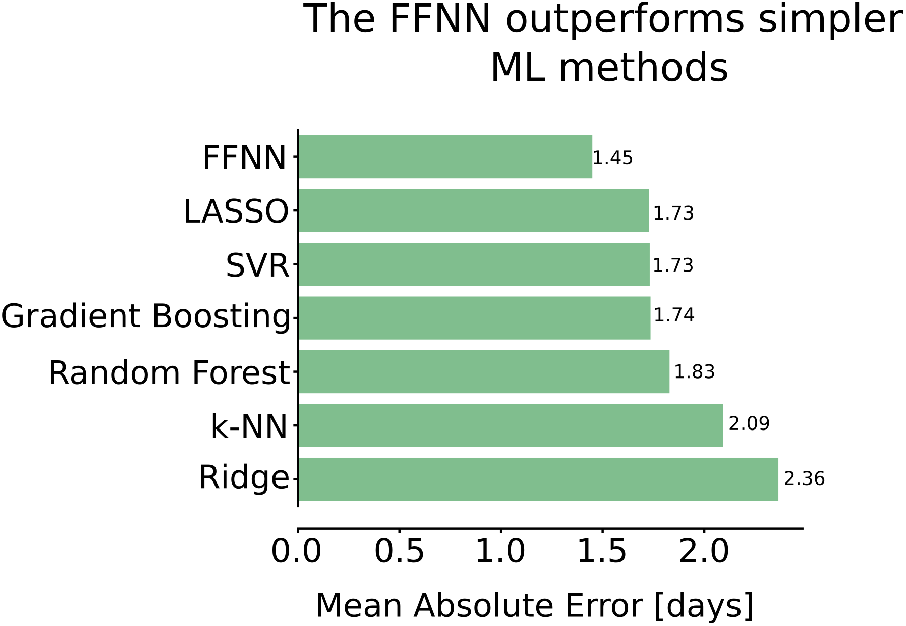
The comparative performance of alternative machine learning methods. We applied a compendium of six supervised regression models to the same metabolomic profiles used to train the FFNN. By evaluating all models on the same test data, we found an ability to predict the PMI across all methods, with the FFNN as the top performer. We here show the MAE of each implemented regression model when applied to this test data.

### Pseudo-time clustering reveals biologically expected metabolomic changes

Having found that all tested machine learning methods satisfactory predicted the time since death from the metabolomic profile, we sought to characterize these post-mortem changes. To this end, we extracted the metabolite features used by the FFNN by selecting the metabolite features for which the activation of the input variable was significantly correlated with the recorded PMI, using a Benjamini-Hochberg corrected p-value of a Spearman’s rank correlation. For these variables (n=810), we created pseudotime series by averaging the abundance of each metabolite feature over individuals with the same PMI. We performed a hierarchical clustering to extract pseudo-time-related groups of metabolites and found three groups of post-mortem changes (Fig. 3). This approach revealed a cluster of 158 metabolite features with decreasing abundance over time, a cluster of 254 metabolite features where the abundance increased, and a cluster of 398 metabolite features where the abundance showed more complex patterns over the pseudo-time scale.

**Figure 3:**
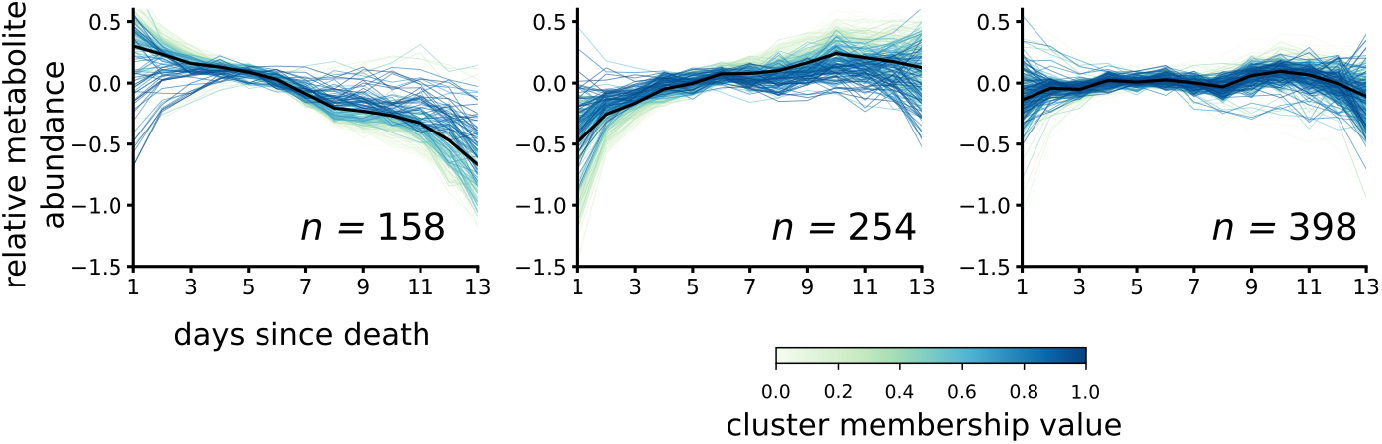
Clustered pseudo-time series reveal broad trends in post-mortem metabolite changes. We computed the pseudo-time series of the metabolites used by the FFNN and divided them into three groups using hierarchical clustering. We found a set with a downward trend containing 158 metabolites, one with an upward trend (254 metabolites), and one with a fluctuating abundance (398 metabolites)

From the visualization of the pseudo-time series in clusters, the more dispersed signal at early (*<*3 days) and late (*>*10 days) PMI becomes visible (Fig. 3). This aligns with the model’s less acurate ability to predict low and high PMI (Fig. 1d). While it is not surprising that longer PMI introduce more uncertainty (e.g. due to surrounding temperatures and other external factors), we wanted to investigate the predictive power of the models at the earlier time points. Therefore, we analyzed the respective predictions of the test data at PMI = 1. We found that all tested models overestimated the actual PMI at day 1, with overestimates ranging from 1.79 (FFNN) to 3.46 (K-NN) days and a median overestimate of 2.56 days (Supplementary Figure S1).

Metabolomic features putatively identified by mass-to-charge matching during functional analysis (MetaboAnalyst) and/or database matching (HMDB) are presented in accordance with cluster membership and relationship to PMI pseudo-time. For features decreasing in relation to PMI pseudo-time, 38 features were identified to 24 unique metabolites (Supplemental Table S2). Of these, acylcarnitines (n = 12) and lysophosphatidylcholines (n = 11) were the most prevalent. For features increasing in relation to PMI pseudo-time 43 features were identified to 30 unique metabolites (Supplemental Table S3). Here, amino acids and dipeptides were the most prevalent. For features fluctuating over PMI pseudo-time, 38 features were identified to 32 unique metabolites (Supplemental Table S4).

### A few hundred samples can be sufficient to accurately predict PMI

Next, we asked how many samples an independent institution needs to train a well-performing model. To answer this, we analyzed how the number of samples/metabolomic profiles in the training dataset influenced model performance. We chose to evaluate the predictive error of a LASSO model as a function of training sample size. Specifically, we randomly selected *n* metabolomic profiles from the training data, with *n*=[16, 32, …, 1024] 150 times for each *n*, and performed a 5-fold cross-validation for model selection. The LASSO regularization parameter *λ* was chosen to minimize the prediction error in a 5-fold cross-validation. Lastly, we evaluated model performance using the independent test dataset.

Interestingly, we found that as few as 256 samples were sufficient to give indications of PMI predictions, achieving an average MAE of 2.05 days (Figure 4). As expected, the prediction error decreased consistently as the number of profiles in the training data increased, underscoring how additional data improved accuracy.

**Figure 4:**
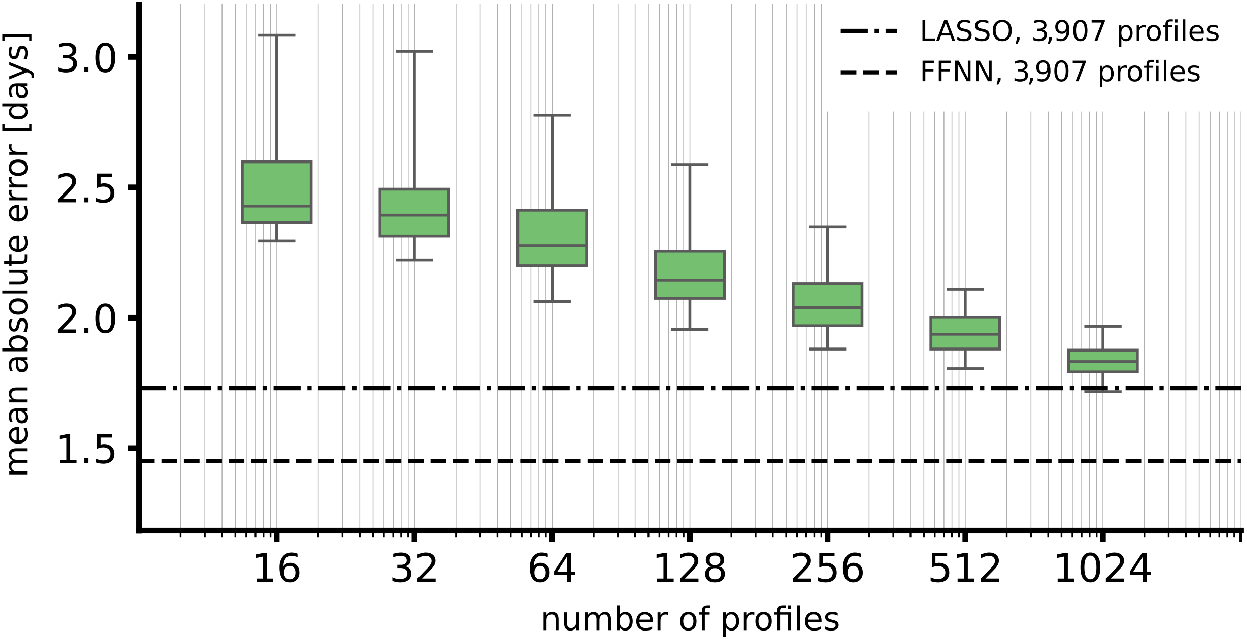
Predictive error as a function of the number of metabolomic profiles used as training data. We randomly sampled n metabolomic profiles from the training data, as shown on the x-axis, to train LASSO models. The y-axis shows the range of the mean absolute error (MAE) of the corresponding 150 models in each set of profiles. The whiskers show the 95% distribution of the respective prediction errors. We found the mean absolute error of this prediction to decrease towards the error of the LASSO model where all 3,907 profiles in the training dataset were used (dashed-dotted line). Shown is also the performance of the full FFNN model on the test dataset (dashed line).

## Discussion

Determining the time of death is one of the central objectives in police and forensic investigations. A PMI estimation can be crucial in crime investigations and court rulings, as well as an important factor in providing closure to next of kin. Despite intensive research, the identification of PMI markers has been slow, and body cooling has remained the gold standard for determining the time since death. Although animal models have demonstrated the potential of metabolomic markers to estimate PMI [8], [13], [14], their application beyond controlled laboratory settings has remained largely untested. Recent research has shown promising results in the use of metabolomic profiling of human body fluids, such as serum, aqueous humor, and vitreous humor to estimate PMI [11], [13], [15]. However, no large-scale human studies have tested the accuracy of the metabolomic data in PMI predictions.

This study demonstrates that the human metabolome paired with machine learning regression models can accurately predict PMI with a mean absolute error of 1.45 days. The error is in line with the uncertainty of the recorded time since death in our collected data. Notably, the uncertainty of the data in our study is 24-48 hours, which means that we have a higher percentage of uncertainty for lower PMI that for higher PMI. Thus, for better estimations of early PMI with our method, we would need higher-resolution PMI data.

Importantly, the combination of metabolomics and machine learning fills the gap of appropriate prediction methods in the crucial time window of 2-10 days (Figure 1d), an interval where established markers often struggle. This finding makes our approach a promising technique for implementation in forensic institutions. We show that the estimations are robust with respect to different machine learning methods, arguably due to the broad signal embedded in post-mortem metabolomic changes. We also visualize the dynamic behavior of the metabolite features by calculating pseudo-time series of each feature across the 4,876 post-mortem metabolomic profiles and cluster features with similar dynamics.

Beyond the early postmortem period, potassium in vitreous humor can be used to estimate PMI. However, predicted PMI from such measurements deviates markedly from the actual PMI after 4-5 days post-mortem [5]. To compare our results with PMI predictions from potassium levels, we used an application developed for this purpose [5]. We assumed ordinary conditions (ambient temperature 20°C and the age of the deceased being 40 years) and estimated PMI with a 95% prediction interval. For potassium concentrations reflecting the early postmortem period (e.g. 10 and 15 mM/L), the application provide PMI estimations of 0.07-1.27 days and 0.08-2.78 days respectively. For potassium concentrations reflecting longer PMI (e.g. 20 and 25 nM/L), the application provide PMI estimations of 0-5.05 days and 0-8.83 days, i.e. broad intervals not useful in forensic practice. These intervals can be compared to our FFNN model with estimated PMI 2.5-5.9 days at PMI=3, and 4.2-8.3 days at PMI=7 (95% intervals). The developed method based on post-mortem metabolomics can thus provide narrower estimations than potassium-based methods in the same range of PMI.

Although we have collected a uniquely large data set of 4,876 metabolomic samples, we also show that substantially lower sample sizes can be used to establish a predictive model for PMI. A sample size as low as 256 used together with a LASSO regression model produced a MAE of 1.73 days, with the error approximately following a linear decrease with exponentially larger samples. We believe that this finding illustrates the broad utility and potential in the development of PMI-predictive models across forensic institutions on a global level.

A great deal of research has been devoted to providing novel methods to predict PMI, often using AI and machine learning methods. Some notable examples include Dai et al. (2019) [8], which trained an SVR on 39 metabolites from 36 Sprague-Dawley rats and reported a mean squared error of 10.33h for PMIs up to 72 h when applied to test data. In a notable paper by Bonicelli et al. (2022) [16], samples of metabolomics, lipidomics and proteomics from human bone samples were used to estimate PMI at 200-800 days, illustrating the long-term relevance. Ferreira et al. (2018) presented how changes in human gene expression could be used to predict PMI up to the first 1,739 minutes, predictive genes being inconsistent between tissues [17].

The post-mortem metabolome is dynamic and changes occur, and according to our data these changes can be used to predict the PMI. This is in line with a recent study by Steuer et al. [12], which evaluated paired samples from the same individuals at two different time points after death. Some of the metabolites in this study change more and some less over PMI, in line with our results (cf. Figure 3). The change in abundance of a certain metabolites with PMI needs to be considered in relation to the change of abundance of the same metabolite as a result of the cause of death in e.g. forensic cause-of-death estimations.

The pseudo-time series in our study (Figure 3) shows that the dynamic features of metabolites can be rather linear over time, possibly allowing for back-calculation to the metabolome at time of death for forensic investigations. This would provide a more accurate description of the metabolome at the time of death, representing a major step forward for forensic investigations of cause-of-death. Furthermore, the success of the linear methods SVR and LASSO to predict PMI (cf. Figure 2), albeit lower than that of the FFNN, indicates that the metabolomic changes as a function of PMI are relatively linear. The demonstration that a LASSO model trained on a subset of metabolomic profiles can adequately predict the PMI (cf. Figure 4) is in line with this observation. It is non-trivial to answer why the performance of the FFNN surpasses that of the other methods, but could be due to mild non-linear dependencies between the metabolomic changes and PMI, Moreover, models such as the LASSO assumes additive effects, while the FFNN could potentially capture combinatorial dependencies between metabolites.

Looking ahead, our data show that post-mortem metabolomics is a possible way forward to get accurate models for PMI prediction. Further validation of our approach in diverse forensic settings is of course essential to ensure its generalizability. We have ongoing collaborations with forensic institutions in other countries and they will be crucial in collecting more varied data, particularly from different environmental conditions and storage practices. Another aspect of the future use of our approach is the development of user-friendly software tools that can be easily adopted by forensic practitioners. This will be a key step in translating our research into practical applications. Ultimately, our goal is to establish a standardized, reliable method for PMI estimation that can be widely implemented, improving the accuracy and efficiency of forensic investigations worldwide.

## Methods

### Study population selection

All autopsy cases admitted between 2017-09-01 and 2019-03-14 at the Swedish National board of forensic chemistry with femoral blood available, ≥18 years of age and that underwent a toxicological screening using high resolution mass-spectrometry were included in the study period (n=7,598). From the study period, all autopsy cases with a confirmed death date or a death date with the same date or at most one-day difference between “body found” and “last seen alive”, were included (in total 4,876 autopsy cases). The most frequent causes of death in the studied population, each with more than 100 documented cases, included complications of cardiovascular disease (n= 748), acute poisoning with one or more drugs (n=659), hanging (n=572), alcohol poisoning (221), drowning (194) trauma resulting in multiple internal and external injuries (150) and gunshot wounds (119). The demographic characteristics of the selected cohort can be summarized as follows: age, median = 56 years (interquartile range = 39-69); sex, male = 3544 (72.3%); BMI, median = 25.7 (interquartile range = 22.5-29.6); PMI, median = 5 days (interquartile range 4-7).

### Calculating the post-mortem interval

In the database of the National Board of Forensic Medicine in Sweden the date of death can be ascribed and coded in two different ways: certain and uncertain. If the date of death is known with certainty this is indicated as such in the database. However, in the majority of cases, the date of death is uncertain. In these cases, a probable date of death is indicated together with the date the deceased was last seen alive. The present study includes all cases in which the date of death is certain (n = 2,876). Additionally, cases with a probable date of death are included if the date when the person was last seen alive is the same or the day before the body was found (n = 2,000). This approach allows for the inclusion of cases where, for example, a person was last seen alive in the evening and found dead the following morning. In some cases, the date of death was ascribed as probable even if the date last seen alive was the same date as when the deceased was found dead. In Swedish forensic practice this is sometimes done to indicate that the death was unwitnessed (i.e. last seen alive in the morning and found dead in the evening).

In the present study PMI is defined as the time (in days) between date of death, certain or probable as described above, to the date of the autopsy in which sampling was performed. Thus, the calculated PMI has a built-in uncertainty of *<*48 hours due to the inclusion criteria above.

### Data processing

The postmortem metabolomic data from the samples were pre-processed in R using the xcms package and CAMERA package, as in [18]. This method extracted 2,305 peaks/metabolic features.

We normalized the data using a log-transform (Eq. 1).

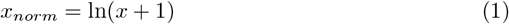

In (Eq. 1), *x* is the peak intensity, and *x*_*norm*_ is the normalized expression. Furthermore, we subsequently standardized the values of each metabolite using a Z-transform. In other words, we subtracted the mean and divided by the standard deviation from each log-transformed metabolomic profile.

We divided the data into training, test, and validation sets in ratios of 80%, 10%, and 10%, respectively. We did this division by randomly assigning each observation to the sets using the respective probabilities.

### Neural network design and training

To predict PMI, we implemented a feed forward neural network regression model. To determine the optimal design of the model, we initiated a hyper-parameter optimization to select the number of hidden layers, the number of hidden nodes in each layer, and the dropout rate for each layer. Additionally, we implemented the option to train the model with a custom attention layer as an input layer such that each metabolite would be individually passed to a single node. The rationale behind this was that such an attention layer would serve as a data-driven feature selection algorithm.

We trained each hyper-parameter setting three times and selected the best performance using an early stopping algorithm with a patience of 25 epochs with respect to prediction error on the validation data. The early stopping algorithm was also used to restore the best weights during training. All tested hyperparameter sets can be found in the Supplementary Table (S1).

### Training of alternative machine learning models

We trained a compendium of alternative machine learning regression models using the same training data as for the FFNN model. These models included the two linear regression models Ridge and LASSO, where the respective *L*_2_ and *L*_1_ penalty weights were selected using the Scikit-Learn build-in RidgeCV and LassoCV implementations. We also implemented a gradient boosting and a random forest regression model, each with the default 100 estimators, as per the default setting on the Scikit-Learn package in Python. Furthermore, we also implemented a K-nearest neighbor (K-NN) regressor and a support vector regression (SVR), both as implemented in Scikit-Learn. We trained the models using the assigned training data, and, for consistency, tested them on the test data.

### Extracting learned model structures and feature identification

We sought to extract which metabolomic features were used in the decision-making processes of the respective models. While such an extraction of input-output dependencies is non-trivial for neural networks, we utilized the attention layer of the FFNN to estimate the usage of input variables. By analyzing these input node activations and correlating them with PMI, we generated a list of metabolomic features ranked by importance.

All metabolomics features included in the FFNN were uploaded to MetaboAnalyst (version 6.0) for functional analysis, which is suitable for untargeted metabolomics data, relying on the assumption that putative annotation at the compound level can collectively predict group level functional changes defined by set of pathways of metabolites [19].

Metabolomic features significant for decision-making in the FFNN were putatively identified and annotated by reviewing the compound hits from the functional analysis in MetaboAnalyst, and/or by matching the mass-to-charge ratio (m/z; *±*5 ppm) to the online human metabolomic database (HMDB; https://hmdb.ca).

## Acknowledgments

The study was supported by the Swedish Research Council (grant no.: Dnr 2023-01407 Green, Dnr 2019-03767 Nyman), the Swedish Fund for Research Without Animal Experiments (grant no.: S2021-0008, F2022-02 Nyman), and the Strategic Research Area in Forensic Sciences at Linkoping University (Magnusson, Ward).

## Author contributions

R.M, C.S., L.J.W, F.C.K, A.E., H.G., and E.N. concieved and planned the experiments. C.S., L.J.W. and A.E. collected and preprocessed the data. R.M., L.J.W., and J.A. performed calculation and modeling analyses with input from all the authors. H.G. and E.N. supervised the work. All authors discussed the results and contributed to the final manuscript.

## Notes

### Competing Interest Statement

The authors have declared no competing interest.

